# Is it reasonable to use of a kinship matrix for best linear unbiased prediction?

**DOI:** 10.1101/568782

**Authors:** Bongsong Kim

## Abstract

The linear mixed model (LMM) is characterized to account for the variance-covariance among entities in a population toward calculating the best linear unbiased prediction (BLUP). Animal and plant breeders widely use the LMM because it is perceived that the a BLUP estimate informs an estimated breeding value (EBV), so to speak a combining ability as a parent, obtained by relating each entity to his/her relatives using the variance-covariance. The LMM practice routinely substitutes an external kinship matrix for the variance-covariance. The challenge relevant to the LMM practice is the fact that it is unrealistic to validate the EBVs because the real breeding values are not measurable but conceptual. This unreality actually means that the EBVs are vague. Although some previous studies measured correlations between the EBVs and empirical combining abilities, they are not sufficient to remove the vagueness of EBVs because uncontrollable environmental factors might interfere with phenotypic observations for measuring the combining abilities. To overcome the challenge, this study scrutinized the soundness of the routine LMM practice from the mathematical perspective. As a result, it was demonstrated that the BLUP estimates resulting from the routine LMM practice mislead the breeding values. The genuine BLUP represents the arithmetic means of multiple phenotypic observations per each entity, given all phenotypic observations adjusted to the mean of zero.

## Introduction

A breeding value means a combining ability of an entity as a parent. The linear mixed model (LMM) is a statistical method that is widely used for estimating breeding values (Piepho, 1994; Panter and Allen, 1995a; Panter and Allen, 1995b; Choi et al, 2017). The fixed-effect variable and random-effect variable are two parameters of LMM. Of these, the random-effect variable has its own variance-covariance, which characterizes the LMM differently from the linear model (linear regression) that is restricted to estimating the fixed-effect variable. The resultant random-effect variable contains estimated breeding values (EBVs) that the variance-covariance implies. It is generally perceived that the variance-covariance in the LMM corrects the breeding-value underestimations and overestimations by accounting for the relationship between entities. Therefore, the resultant random-effect variable is termed best linear unbiased prediction (BLUP) (Henderson, 1975).

Routine LMM practices substitute an external kinship matrix resized toward obtaining the least square for the variance-covariance (Robinson, 1991). A number of previous studies validated the effectiveness of BLUP taking advantage of the kinship matrix by empirically measuring the combining abilities through field tests and computer simulations (Belonsky and Kennedy, 1988; Piepho, 1994; Panter and Allen, 1995a; Panter and Allen, 1995b; Bauer et al, 2006; Nielsen et al, 2011; Choi et al, 2017; Manzanila-Pech et al, 2017). However, the empirical validations necessarily limit thorough evaluation of the quality of EBVs because it is impossible to obtain exact combining abilities to be compared with the EBVs. To overcome the limitation, this study investigated the integrity of BLUP from the mathematical perspective.

## Materials and Methods

### Naive-BLUP

The LMM is denoted as:

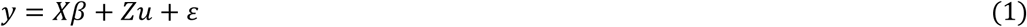

where *y* is the phenotypic variable; *X* is the design matrix for fixed effect; *Z* = the design matrix for random effect; *β* = the unknown fixed-effect variable; *u* = the unknown random-effect variable; *ε* = the unknown error-term variable.

Equation 1 assumes *ε*~*N*(0,*Iσ*^2^). The *u* contains EBVs and can be calculated as follows:

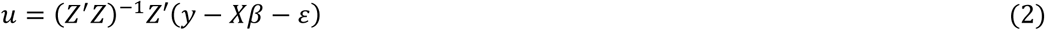

The *var*(*u*) can be calculated as follows:

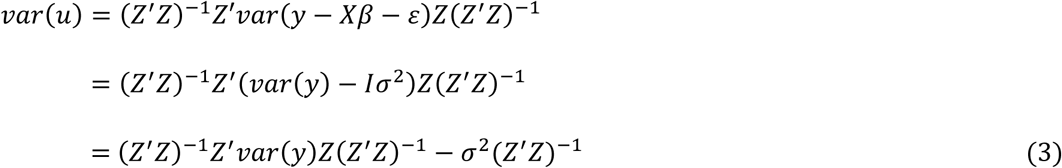

The *var*(*y*) can calculated as follows:

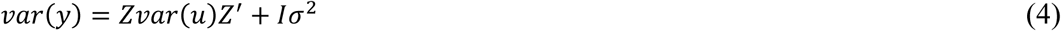

Let us substitute Equation 4 for Equation 3 as follows:

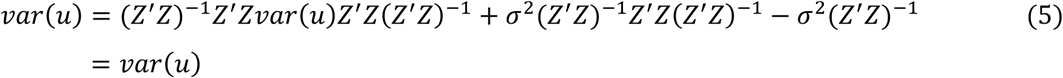

Equation 5 demonstrates that the *σ*^2^(*Z′Z*)^−1^ in Equation 3 is negligible because it will be cancelled by *var*(*y*). For simplicity, therefore, Equation 3 can omit the *σ*^2^(*Z′Z*)^−1^ so that:

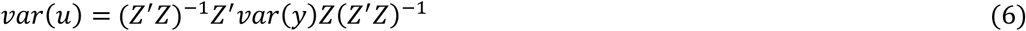

Because 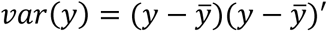, Equation 6 can be written as follows:

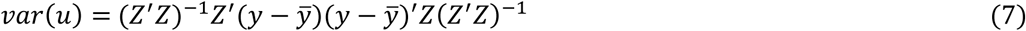

where 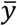 is a vector containing the averages of *y*.

Equation 7 is equivalent to the following equation:

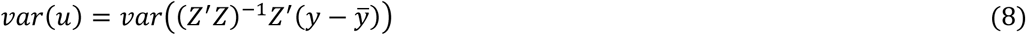

Therefore,

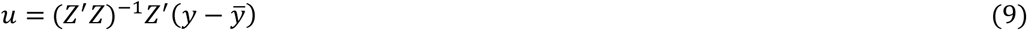

Equation 9 is derived from statistical basics. Hereafter, the *u* resulting from Equation 9 is called Naive-BLUP. Every value of *u* for Equation 9 represents the arithmetic mean of multiple phenotypic observations per each entity, given all phenotypic observations adjusted to the mean of zero. Once *u* is known, *β* for Equation 1 can be calculated using the following equation:

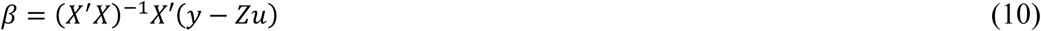

### Average-vector

If the Naive-BLUP does not adjust all phenotypic observations to the mean of zero, the resultant vector will contain the arithmetic means of multiple phenotypic observations per each entity, which can be calculated as follows:

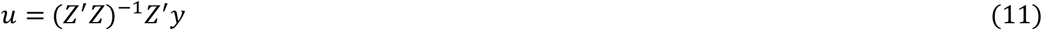

Hereafter, the *u* resulting from Equation 11 is called Average-vector.

### K-BLUP

The equation for the routine LMM is as follows:

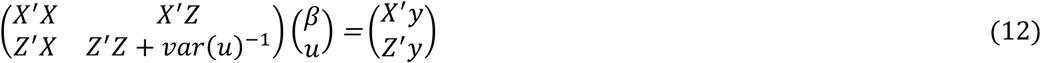

Equation 12 was originally derived by Henderson (1975). The exact year of its first release is unclear. The mathematical derivation of Equation 12 from Equation 1 is available in a lecture note by Searle (1980).

In Equation 12, the *u* can be obtained using the row operation as follows (Kim et al. 2018):

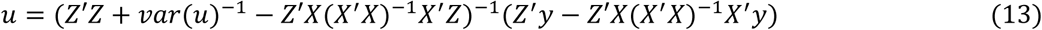

If *var*(*u*) obtained by Equation 7 is used, Equation 13 always produces the same result as Equation 9. However, the routine LMM practice assumes that *var*(*u*) = *λK*, where *K* is the kinship matrix; 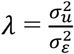, where 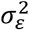 and 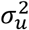 are the variance components for *ε* and *u*, respectively. Therefore, Equation 13 can be transformed into the following form:

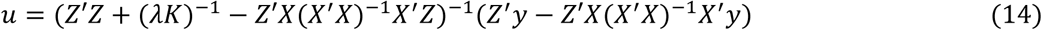

Equation 14 comprises the *K* (kinship matrix). Hereafter, the u resulting from Equation 14 is called K-BLUP. Equation 10 can be commonly used for calculating the *β*. This study fitted the K-BLUP using the expectation-maximization (EM) algorithm using GWASpro (Kim et al, 2018). More details about the EM algorithm can be found in the related paper.

### Rice data set

To compare between two *u*’s obtained by the Naive-BLUP and K-BLUP, an open-access rice data set comprising phenotypic and SNP (single nucleotide polymorphism) data was used, which was downloaded from http://ricediversity.org/data/index.cfm. The data set was originally prepared, analyzed and released by Spindel et al. (2015). Therefore, more details can be found in the related paper. The phenotypic trait used in this study was grain yield (kg/ha), which had the structure of three dimensions: four years (2009, 2010, 2011, 2012), two seasons (DS: dry season; WS: wet season) per each year and three replications (1, 2, 3) per each season. The three dimensions were reduced to two by averaging the three replications. Consequently, the yield observations were arranged into eight environments: 2009/DS, 2009/WS, 2010/DS, 2010/WS, 2011/DS, 2011/WS, 2012/DS and 2012/WS. Originally, 359 rice accessions were present in the phenotypic data. Ultimately, 107 rice accessions remained after deleting accessions which had missing phenotypic observations or were absent in the SNP data. Accordingly, the SNP data set was aligned with the phenotypic data, which resulted in 107 accessions genotyped with 108,024 SNPs. The SNP data set was used to calculate a *K* using Numericware i (Kim and Beavis, 2017).

### Validating the integrity of K-BLUP

To examine the integrity of K-BLUP, the following proposition was established:

**Proposition 1.** The integrity of K-BLUP is secured, only if 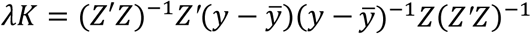.

**(Proof)** The K-BLUP defines *var*(*u*) = *λK*. The process from Equations 1 to 7 shows that the *var*(*u*)in the LMM must hold the property of Equation 7 which is 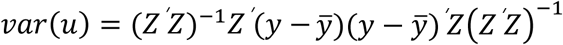. In order for the K-BLUP to be correct, therefore, it must be satisfied that 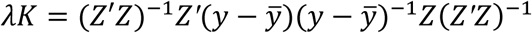.

Meanwhile, the Naive-BLUP is based on Equation 7 so that the property of Equation 7 is naturally held. To compare the two *var*(*u*)s calculated by *λK* and Equation 7, the Pearson correlation coefficient between their lower triangle matrices was calculated.

### Repository of input data and computer code

All input data and computer code are freely available at https://github.com/bongsongkim/BLUP.

## Results

### Comparison between Naive-BLUP, K-BLUP and Average-vector

The Naive-BLUP, K-BLUP and Average-vector were calculated using the given rice data set. Figure 1A shows that the correlation coefficient between the resultant K-BLUP and Naive-BLUP was 0.9526. Figure 1B shows that the correlation coefficient between the resultant Naive-BLUP and Average-vector was 1.0, illustrating the complete proportionality between them. Figure 1C shows that the correlation coefficient (0.9526) between the resultant K-BLUP and Average-vector was the same as the case of Figure 1A, which is due to the complete proportionality between the Naive-BLUP and Average-vector. Table 1 summarizes the resultant K-BLUP, Naive-BLUP and Average-vector and shows that the grand mean of Naive-BLUP is zero. Figure 1B and the mean of zero indicate that every value for Naive-BLUP represents the average of multiple phenotypic observations per each entity, given all phenotypic observations adjusted to the mean of zero.

**Table 1.** Summaries of the resultant K-BLUP, Naive-BLUP and Average-vector.

|  | MI n. | 1<sup>st</sup> Qu. | Medi an | Mean | 3<sup>rd</sup> Qu. | Max. |
| --- | --- | --- | --- | --- | --- | --- |
| K-BLUP | -253. 58 | 30. 46 | 118. 45 | 113. 18 | 201. 40 | 381. 81 |
| Naï ve-BLUP | -540. 5 | -130. 9 | 18. 8 | 0. 0 | 117. 1 | 403. 8 |
| Average-vector | 2445 | 2855 | 3005 | 2986 | 3103 | 3390 |

**Figure 1.**
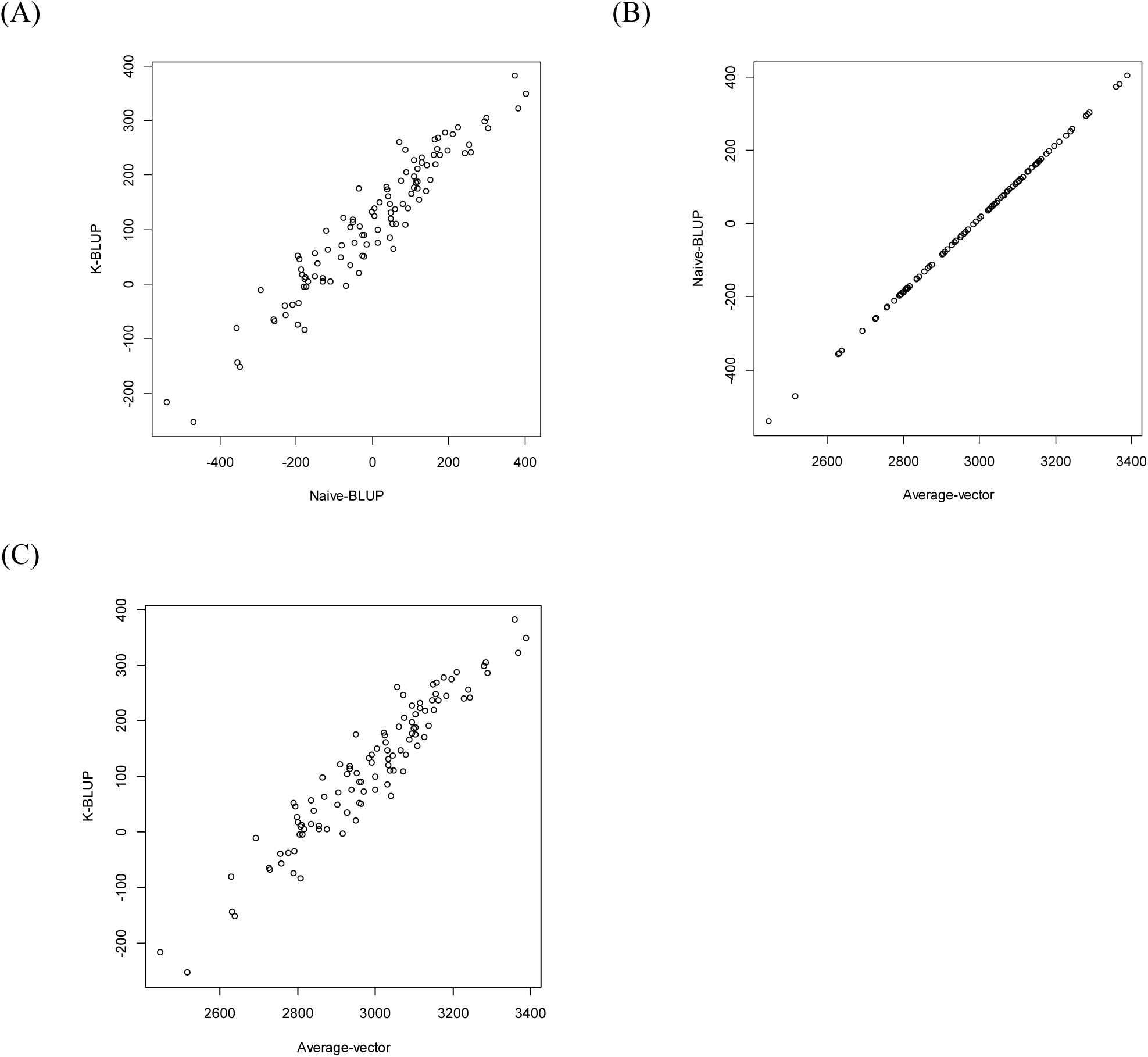
(A) The correlation plot between the resultant K-BLUP and Naive-BLUP. (B) The correlation plot between the resultant Naive-BLUP and Average-vector. (C) The correlation plot between the resultant K-BLUP and Average-vector.

### Comparing *var*(*u*)s resulting from K-BLUP and Naive-BLUP

Two *var*(*u*)s resulting from the Naive-BLUP and K-BLUP were computed using the same rice data set. Sub-matrices of the former and the latter are shown in Tables 2 and 3, respectively. The *var*(*u*) resulting from the Naive-BLUP was calculated using Equation 7, while the *var*(*u*) resulting from the K-BLUP was calculated by multiplying the *K* by *λ*. The *λ* is the constant that leads to the least square of LMM, which can be calculated by 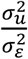, where 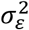 and 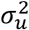 are the variance components for *ε* and *u*, respectively. The resultant estimates for 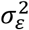 and 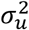, obtained using the EM algorithm, were 80019.630 and 86076.655, respectively, which resulted in *λ* = 1.076. Therefore, Table 3 was obtained by multiplying the *K* by 1.076. The comparison between Tables 2 and 3 showed the two *var*(*u*)s are different, leading to the conclusion that Proposition 1 is not satisfied. Figure 2 shows the correlation plot between two lower triangle matrices of *var*(*u*)s calculated by *λ*K and Equation 7. The resultant correlation coefficient was 0.099, indicating that the two *var*(*u*)s were hardly correlated.

**Table 2.** The sub-matrix of *var*(*u*) obtained by Equation 7 using the rice data set.

|  | A1257 | A1258 | A1259 | A1260 | A1261 | A1262 | A1263 | A1264 | A1265 | A1267 |
| --- | --- | --- | --- | --- | --- | --- | --- | --- | --- | --- |
| A1257 | 2608.254 | -13200.4 | 3938.23 | -5769.53 | -2119.01 | -19106.5 | -5967.43 | -10053.1 | -2454.17 | -6556.87 |
| A1258 | -13200.4 | 66807.05 | -19931.4 | 29199.58 | 10724.32 | 96698.1 | 30201.15 | 50878.8 | 12420.53 | 33184.33 |
| A1259 | 3938.23 | -19931.4 | 5946.374 | -8711.47 | -3199.52 | -28849.1 | -9010.28 | -15179.3 | -3705.57 | -9900.29 |
| A1260 | -5769.53 | 29199.58 | -8711.47 | 12762.36 | 4687.312 | 42264.16 | 13200.12 | 22237.77 | 5428.682 | 14503.99 |
| A1261 | -2119.01 | 10724.32 | -3199.52 | 4687.312 | 1721.539 | 15522.63 | 4848.091 | 8167.406 | 1993.827 | 5326.972 |
| A1262 | -19106.5 | 96698.1 | -28849.1 | 42264.16 | 15522.63 | 139963.1 | 43713.86 | 73643.17 | 17977.77 | 48031.79 |
| A1263 | -5967.43 | 30201.15 | -9010.28 | 13200.12 | 4848.091 | 43713.86 | 13652.89 | 23000.54 | 5614.891 | 15001.49 |
| A1264 | -10053.1 | 50878.8 | -15179.3 | 22237.77 | 8167.406 | 73643.17 | 23000.54 | 38748.19 | 9459.205 | 25272.47 |
| A1265 | -2454.17 | 12420.53 | -3705.57 | 5428.682 | 1993.827 | 17977.77 | 5614.891 | 9459.205 | 2309.18 | 6169.513 |
| A1267 | -6556.87 | 33184.33 | -9900.29 | 14503.99 | 5326.972 | 48031.79 | 15001.49 | 25272.47 | 6169.513 | 16483.29 |

**Table 3.** The sub-matrix of *var*(*u*) obtained by *λK* using the rice data set.

|  | A1257 | A1258 | A1259 | A1260 | A1261 | A1262 | A1263 | A1264 | A1265 | A1267 |
| --- | --- | --- | --- | --- | --- | --- | --- | --- | --- | --- |
| A1257 | 2.151388 | 1.851162 | 1.850721 | 1.800379 | 1.812319 | 1.731394 | 1.827389 | 1.873655 | 1.873053 | 1.885466 |
| A1258 | 1.851162 | 2.151388 | 1.849775 | 1.811587 | 1.779897 | 1.664529 | 1.797334 | 1.865082 | 1.826185 | 1.817923 |
| A1259 | 1.850721 | 1.849775 | 2.151388 | 1.722724 | 1.768743 | 1.662496 | 1.772594 | 1.926719 | 1.78477 | 1.801519 |
| A1260 | 1.800379 | 1.811587 | 1.722724 | 2.151388 | 1.795947 | 1.659366 | 1.8301 | 1.814675 | 1.924923 | 1.874623 |
| A1261 | 1.812319 | 1.779897 | 1.768743 | 1.795947 | 2.151388 | 1.801766 | 1.742151 | 1.792795 | 1.796958 | 1.796097 |
| A1262 | 1.731394 | 1.664529 | 1.662496 | 1.659366 | 1.801766 | 2.151388 | 1.724833 | 1.705642 | 1.719659 | 1.715926 |
| A1263 | 1.827389 | 1.797334 | 1.772594 | 1.8301 | 1.742151 | 1.724833 | 2.151388 | 1.800562 | 1.835156 | 1.846408 |
| A1264 | 1.873655 | 1.865082 | 1.926719 | 1.814675 | 1.792795 | 1.705642 | 1.800562 | 2.151388 | 1.8696 | 1.837092 |
| A1265 | 1.873053 | 1.826185 | 1.78477 | 1.924923 | 1.796958 | 1.719659 | 1.835156 | 1.8696 | 2.151388 | 1.876742 |
| A1267 | 1.885466 | 1.817923 | 1.801519 | 1.874623 | 1.796097 | 1.715926 | 1.846408 | 1.837092 | 1.876742 | 2.151388 |

**Figure 2.**
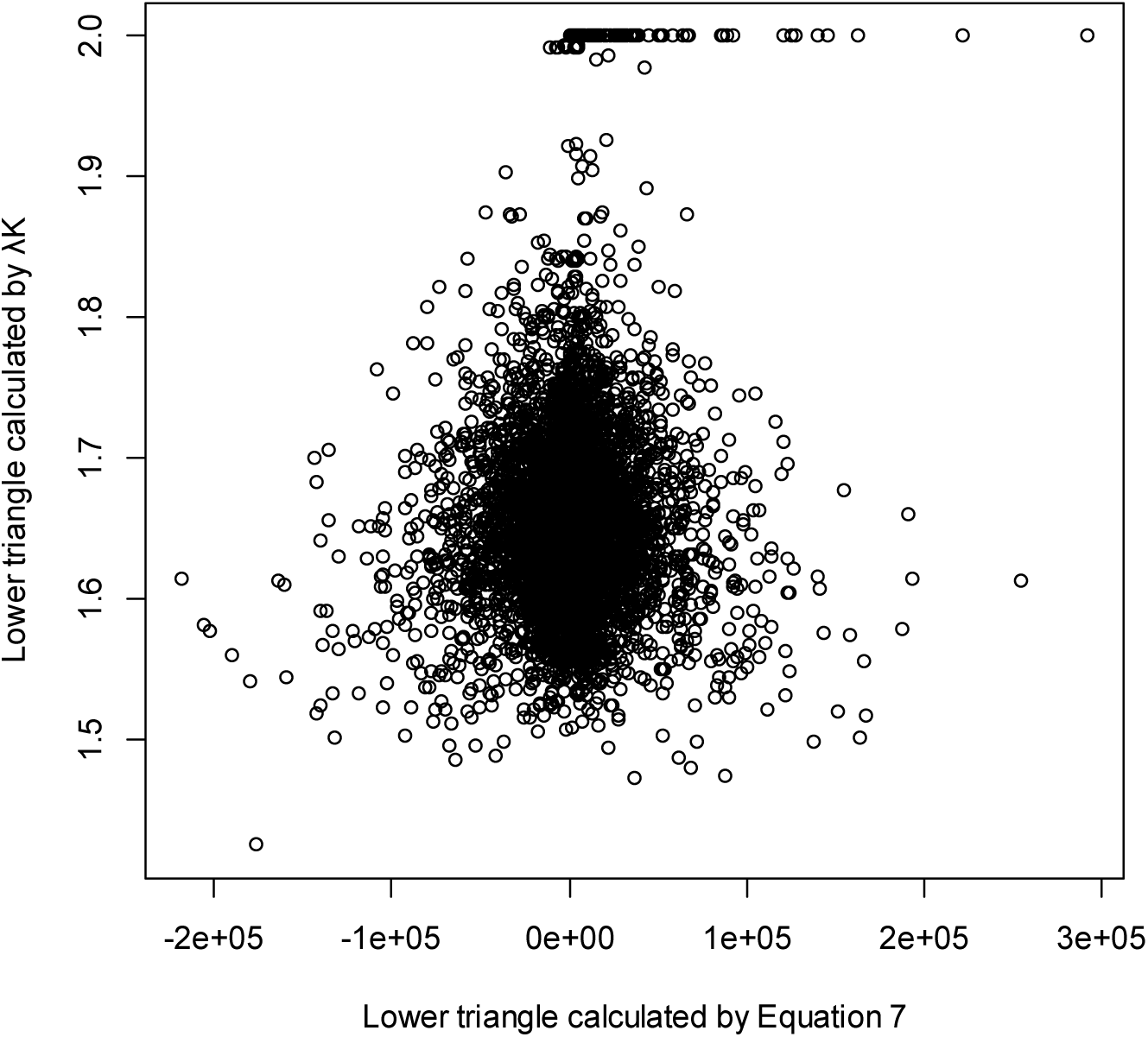
Correlation plot between two lower triangle matrices of *var*(*u*)s calculated by *λK* and Equation 7. The resultant correlation coefficient was 0.099.

## Discussion

The mathematical process from Equations 1 through 9 for the Naive-BLUP shows the whole property of BLUP. Of the process, the relationship between *var*(*u*) and *var*(*y*) in Equation 7 indicates that *u* depends on *y*. This means that the direction of *u* is influenced by *y*, in which it is important to note that the direction of *u* implies the ranking of entities by breeding value. This is a crucial property of BLUP because the phenotypic variable (*y*) must be a key input for the breeding-value variable (*u*). Equation 9 shows that, given all phenotypic observations adjusted to the mean of zero, the Naive-BLUP results in a vector containing the arithmetic means of multiple phenotypic observations per each entity. Figure 1B suggests that the Naive-BLUP is proportional to the Average-vector, illustrating that the latter implies the former. Considering that the average of multiple phenotypic observations can represent how well an entity adapts to multiple environments, its association with the breeding value is reasonable (Piepho, 1994). This is because the meaning of breeding value must comprehend phenotypic variation for an entity. In other words, a single phenotypic observation cannot be eligible for a breeding value because the phenotypic variation is missing.

Meanwhile, the K-BLUP assumes that *var*(*u*) = *λK* other than Equation 7. Considering that the *λ* is a scholar, the direction of *u* is solely determined by *K*. This causes a critical problem in that *y* cannot influence the direction of *u*. This problem illustrates that the assumption of K-BLUP fundamentally has a flaw. In actuality, however, the dissatisfaction of Proposition 1 means that *λK* does not result in *var*(*u*), indicating that the assumption of K-BLUP cannot be satisfied. This doubles the flaw of K-BLUP with regard to the assumption that *var*(*u*) = *λK*. The discrepancy between two *var*(*u*)s resulting from the Naive-BLUP and K-BLUP is shown in the comparison between Tables 2 and 3 as well as Figure 2. Another flaw of K-BLUP is the dilemma that *var*(*u*) must be known to estimate the unknown *u*, which is unrealistic. To overcome this unreality, the routine practice of K-BLUP substitutes *λK* for *var*(*u*) (Yu et al, 2006; Bradbury et al, 2007; Gilmour et al, 2009; Lipka et al, 2012; Kim et al, 2018). However, *λK* merely implies either the genomic similarity based on DNA-sequence homology or genealogical co-ancestry based on pedigrees, other than the variance-covariance among EBVs (Emik and Terrill, 1949; Henderson, 1976; VanRaden, 2008; Kim et al, 2016; Kim and Beavis, 2017). Therefore, the use of *λK* as the substitution for the unknown *var*(*u*) in the K-BLUP misleads the breeding values.

## Conclusion

The *var*(*u*) in the Naive-BLUP soundly secures the integrity of BLUP as well as correct breeding values. The soundness indicates that the Naive-BLUP is the genuine BLUP.

